# The forgotten wine: understanding the ecology and composition of palm wine fermentation

**DOI:** 10.1101/2024.04.26.591403

**Authors:** I Nyoman Sumerta, Xinwei Ruan, Kate Howell

## Abstract

Palm wine is an alcoholic beverage that has existed for centuries and has important economic and socio-culture values in many tropical and sub-tropical countries. Lesser known than other types of wines, palm wine is made by spontaneous fermentation of palm sap by naturally occurring microbial communities. The palm sap ecosystem has unique microbial composition and diversity, which determines the composition of the eventual wine and is likely affected by geographical distinctiveness. While these features are well understood in grape and rice wine, these features have not been understood in palm wine. In this review, we gather information of microbial communities and metabolite profiles from published studies, covering a wide range of methodologies and regions, to better understand the causal links between the principal microbial species and major metabolites of palm wine. We assess palm wine quality across production regions and local practices to provide general characteristics of palm wine and identify specific regional information. These will provide better understandings to the function of microbial communities and metabolite diversity, the contribution of regional variations and to ensure product quality in this important and widespread, yet overlooked, fermented beverage.

**One sentence summary:** review and synthesis of microbial ecology and metabolites in palm wine fermentation across geography and their contribution to cultural food systems

## 1. Introduction

Fermenting microbial communities form beverage aromas and flavours, by release or transformation of chemical compounds in the substrate, creation of small molecules during microbial metabolism and as a result of ecological interactions (Conacher et al., 2021; Liu et al., 2019; Smid & Lacroix, 2013). Other contributing factors to the determination of beverage sensory characteristics are regional variations of plant species and cultivar, genetic diversity of plant and microbes, all of which affect the chemical profile of the end product (Bokulich et al., 2016; Liu et al., 2020). We know that there are a multitude of dynamic and complex microbial and metabolic interactions, which impact upon the product quality of fermented beverages and this has been a major challenge for product development (Gobert et al., 2022). In grape growing and winemaking, understanding and defining the distinctiveness of microbes in fermentation preserves sensory diversity and increases the product value (Bokulich et al., 2016). For example, protecting designations of origins (PDOs) and geographical indication (GIs) of fermented beverages have a significant impact in economic aspect of the product (Candiago et al., 2022; Deselnicu et al., 2013).

Palm wine is less acknowledged compared to the better-known fermented beverages made from grapes and rice. Palm wine is an extremely important commodity in many tropical countries as a traditional beverage, contributing to local economies and with distinctive impacts upon culture and community (Das & Tamang, 2021; Djeni et al., 2020; Santiago-Urbina & Ruíz-Terán, 2014; Wijaya et al., 2024). Palm wine is produced as a traditional practice and is treated differently based on its origin, which leads to distinctive regional variation. In common to all production systems is the collection of palm sap from fruiting bodies and fermentation.

The fermentation of palm wine relies on native microbial communities, which are likely have adapted to local biogeographical constraints to conduct the biotransformations (Astudillo-Melgar et al., 2019). Some microbial species (including yeasts) have evolved to occupy a niche to engage with a wider range of ecological interactions (Gray & Head, 2008). However, biotic and abiotic factors can specifically modify microbial community structure (Zhou et al., 2021), and drive the differentiation of evolutionary traits and chemical release of the principal fermentative species, *Saccharomyces cerevisiae* (Knight et al., 2015). Understanding biogeographical ranges of microbial communities will contribute to the comparison of global populations, especially on domestication trajectory of palm wine’s principal species (De Guidi et al., 2023; Ezeronye & Legras, 2009; Tra Bi et al., 2019) and gain cultural traction by being associated with the regionality of palm wine from topical countries.

Spontaneous fermentation of palm wine demonstrates dynamic nature and changing occurrence of fermentative and non-fermentative microbes, which lead to flavour formation in the finished beverage. Understanding microbial ecology is important to develop a consistent beverage product (Ciani et al., 2012). However, published reports of palm wine ecology, especially covering microbial and metabolite profiles and with broad geographical origins are limited. Gathering and compiling recent studies can construct a general pattern of palm wine fermentation. For example, comprehensive studies of microbial communities and regional characteristics of grape wine have helped understanding of how the fermentation technique and ecological factors influence product specificity (Bokulich et al., 2016; Knight et al., 2015). Examining palm wine microbial ecology across tropical areas will give new insights into the range, activity and formation of yeast and bacterial communities in sap from palm trees.

This review focuses on gathering the principal microbial communities and chemical compounds found during palm sap fermentation, that represent the characteristics of palm wine linked to ecology and geography. We begin by discussing the distinctive composition of palm wine and the geographical areas of palm wine production. We synthesise information from published studies to explain the contribution of ecological factors on the fermentation process and the likely effects on organoleptic properties of palm wine. We conclude by examining the influence of local cultural practices on palm wine, followed by suggestions for directions in product development.

## 2. The distinctive composition and flavour of palm wine

Grapes and rice wines are the most known types of fermented beverages across the world and play an important part in social and economic development of many countries (Anderson & Wittwer, 2015; Bokulich et al., 2016). Meanwhile, palm wine from topical countries is largely unknown. Palm wine is a traditional alcoholic beverage in Africa, south Asia, southeast Asia, and central America (Santiago-Urbina & Ruíz-Terán, 2014). Palm wine is derived from palm sap, which is tapped form the blossom or apical structure of palm tree (Djeni et al., 2020; Hebbar et al., 2018). The fresh sap drips from the cutting part with slightly white turbid colour and moderate sweetness (Chandrasekhar et al., 2012), containing sugars (predominantly sucrose) (Naknean et al., 2010; Limtong et al., 2020). In grapes, fructose and glucose predominate (Corbin et al., 2015) and a crushing step is needed to release these sugars from the grapes to initiate the fermentation process. Rice requires extensive processing before starch conversion is ready to ferment (Lee et al., 2022). In contrast, palm sap can be directly fermented after tapping without any pre-processing steps. After fermentation, palm wine has about 4-6% ethanol by volume (Das & Tamang, 2021; Santiago-Urbina & Ruíz-Terán, 2014), which is relatively similar to sweet rice wine 4.53% (v/v) (Yang et al., 2022). On the other hand, grape wine is higher about 11-15% (v/v) (Fontoin et al., 2008) and has more total phenolics at around 1103 mg/L (Ivanova et al., 2010), compared to palm wine up to 340 mg/L (Das & Tamang, 2023) and 250 mg/L in sweet rice wine (Yang et al., 2022). For total flavonoids content, palm wine has higher value up to 490 mg/L (Das & Tamang, 2023) than red grape wine in 237 mg/L (Ivanova et al., 2010) and 10 mg/L in sweet rice wine (Yang et al., 2022). Despite the variance in these substrates, these plant substrates all support the growth of yeast and bacteria during fermentation.

Palm sap is a versatile commodity from palm trees, that can be transformed into further derivative products (**Fig. 1**). The fresh sap can be directly consumed and has various beneficial metabolites, including organic acids (Doddipalla et al., 2022; Naknean et al., 2010; Nwaiwu et al., 2016; Pammi et al., 2021), vitamins (Barh & Mazumdar, 2008; Zongo et al., 2020), proteins (Pammi et al., 2021), phenolics (Xia et al., 2011), and minerals (Barh & Mazumdar, 2008). A shift from sugar fermentation to acidic fermentation generates a variety of derivative products, such as palm sugar, alcoholic beverages, and palm vinegar (González & De Vuyst, 2009; Limtong et al., 2020; Somawiharja et al., 2018; Zongo et al., 2020). The fresh sap can be refined to obtain palm sugar and left naturally at room temperature to produce palm wine. Palm wine typically has shelf life only for 1-2 days of fermentation for drinking pleasantly before a subsequent acidic fermentation, which change the flavour. Palm wine can be distilled to produce spirits, which has more than 40% ethanol (v/v) (Pratiknjo & Mambo, 2019). A longer fermentation generates organic acids as a source to produce a palm vinegar. In all cases, the typical fermentation is conducted by palm’s natural microbial community without addition of cultured microbes and in general swift and dynamic fermentation process occur (Das & Tamang, 2021; Djeni et al., 2020).

**Fig. 1.**
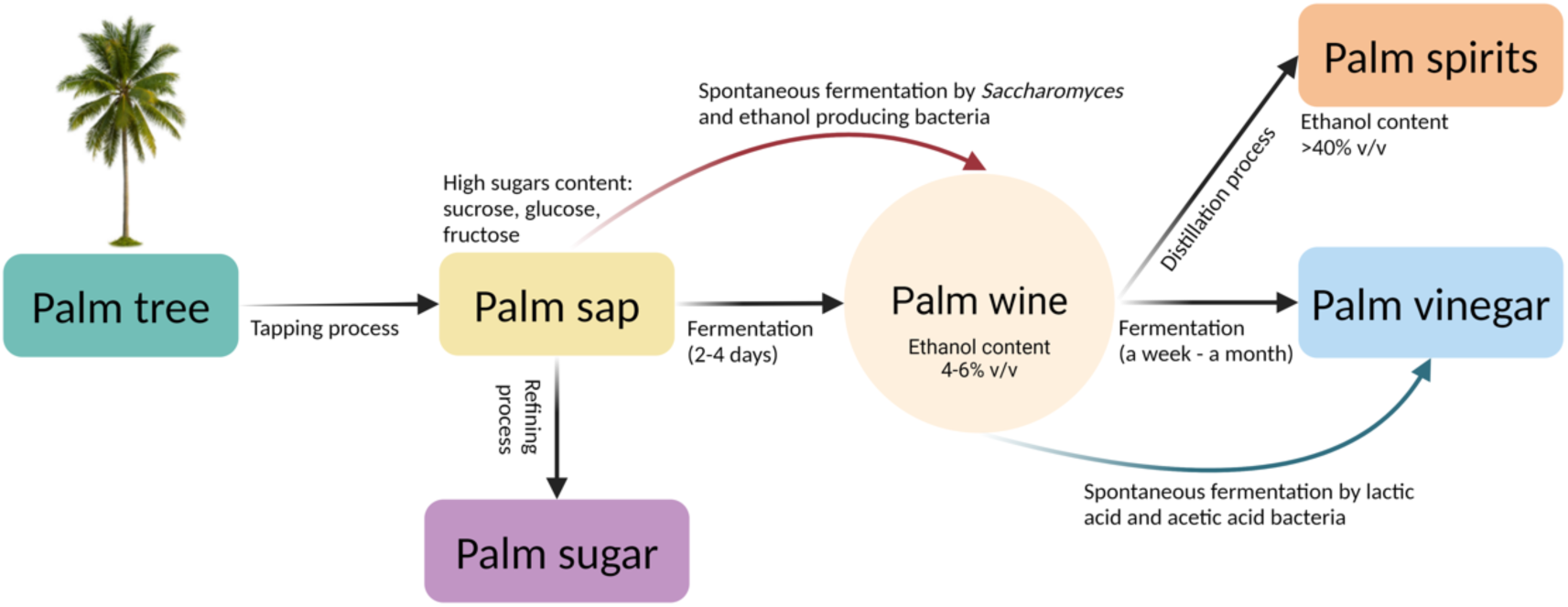
Schematic to describe the processing of palm sap with its derivative products. Created with BioRender.com

## 3. Biogeographical perspectives of palm wine

Palm trees are differently distributed across sub-tropical and tropical countries. The use of specific palm species and the influence of ecological factors may mediate microbial ecology, which determines the distinctiveness of palm wine sensory attributes. We have compiled recent studies to construct a clear picture of palm wine usage and tapping practices. By collated data from 47 studies, specifically focused on microbial and biochemical diversity topics within various approaches (**Supplementary Fig. S1**). Based on our compiled studies, palm wine has been studied across three continents over 14 tropical and subtropical countries (**Fig. 2A**). More than half of the studies have originated in Africa, where reports from production practices in Burkina Faso predominate, particularly in metabolite and microbial studies, followed by India. Each continent uses specific palm species for palm wine making; *Nypha* sp. in Asia and *Acrocomia* sp. in Central America, while *Elaesis* spp. and *Raphia* spp. are specifically found in Africa. The traditional names of palm wine vary across the world. In Burkina Faso and Cote d’Ivoire, palm wine is widely known as *Bandji* (Karamoko et al., 2012; Ouoba et al., 2012); *Mimbo* in Cameroon (Ciani et al., 2012; Jespersen, 2003); *Akpeteshie* in Ghana (Amoa-Awua et al., 2007); *Toddy* in India (Tamang, 2022); *Tuak* in Indonesia (Hermansyah et al., 2015; Wijaya et al., 2024); *Taberna* or *Tavern* in Mexico (Coutiño et al., 2020; Santiago-Urbina & Ruíz-Terán, 2014); *Ogogoro* in Nigeria (Ikeh et al., 2021); *Tuba* in Philippines (Gregorio, 2023); *Ra* in Sri lanka (Ciani et al., 2012); *Songkhla* in Thailand (Naknean et al., 2010); *Lagmi* in Tunisia (Malek et al., 2017).

**Fig. 2.**
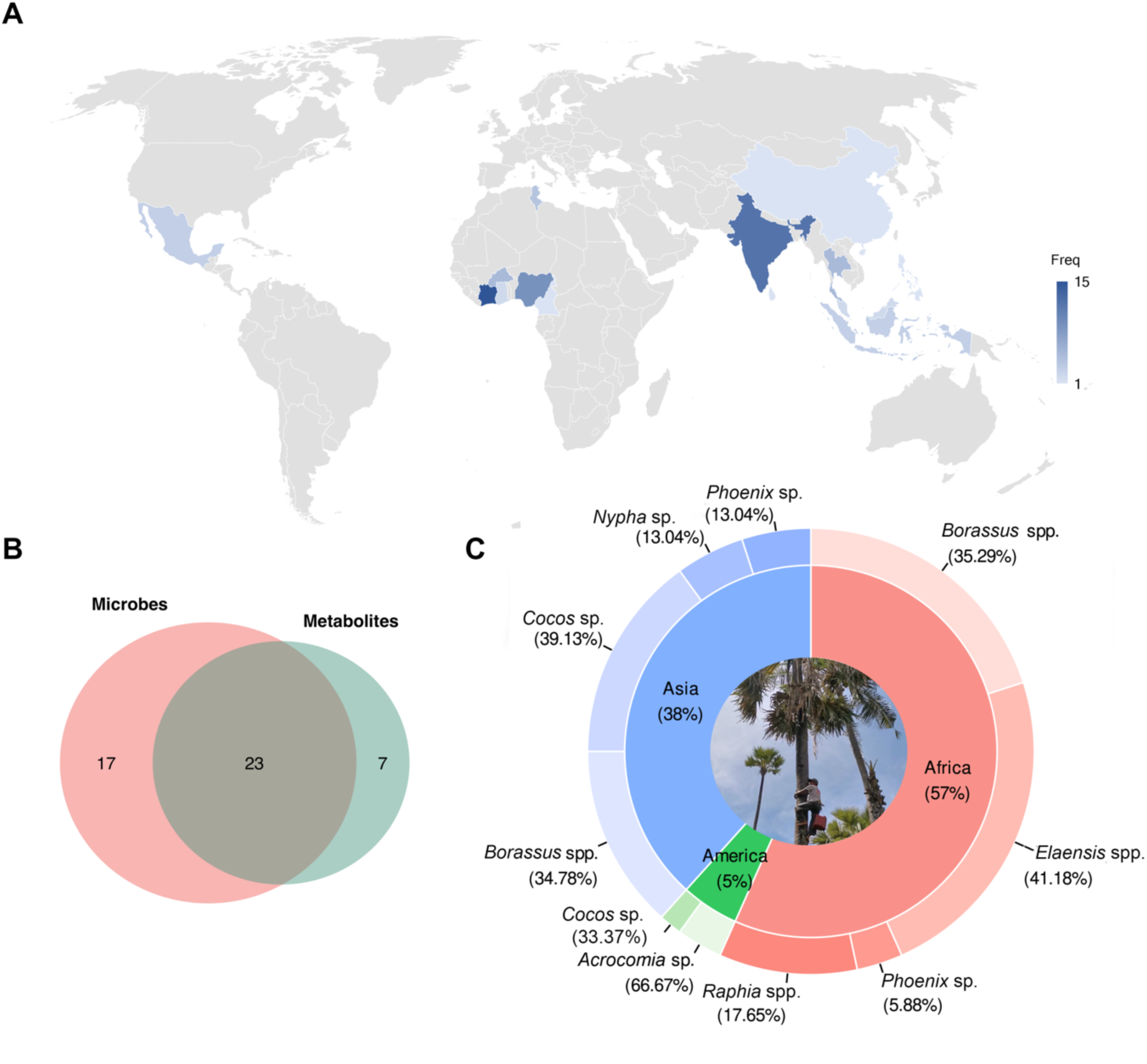
A survey of palm wine studies which include measures of microbial ecology and metabolite profiling (n= 47), details are available on **Supplementary Table S1 & S2**. A) Country distribution of scientific studies in both microbes and metabolites across continents. B) The Venn diagram shows the focus of compiled scientific studies. C) Proportion of the studies of palm species for all compiled studies.

### 3.1 Geography and palm species affects palm wine production

Environmental variables have a significant role on palm sap production and characteristics (Ziadi et al., 2014). Although it may be less appreciated, environmental factors contribute to the palm physiology, which can directly affect the sap yield and physicochemical characters, inducing the microbial communities, and alter the fermentation process. Astudillo-Melgar et al. (2019) believed that microbial diversity is correlated to biogeographical factors from where palm wine is produced and adapted with specific ecological variables. Factors such as climate, weather, palm species, microbial vector, and tapping process may not work independently to alter the microbial composition, but often exert an integrated effect. Therefore, impacts will be discussed as an important aspect to taking further action on expected mitigation for product development.

Although palms are mostly observed in coastal areas of tropical and sub-tropical countries (Moore & Uhl, 2019), the variety of local factors could also impact the diversity of microbial community. Barata et al. (2012) emphasised that microbial diversity is relied on climatic condition, temperature, sunlight exposure, wind, and rainfall where the plant origin, indicating the sign of terroir. Environmental factors have been observed on their influence for plant and soil-microbes association (Zhou et al., 2021); however, there is a lack information on arboreal microbes correlated to palm sap tapping. A study by Ziadi et al. (2014) reported that sap production and physicochemical characters are significantly influenced by tapping season. Higher sugar content and less acidic are characterized from palm sap in warmer seasons, which have more sunlight exposure to increase palm photosynthesis and thus sugar content. Microbial counts are slightly higher, which is less affected by the warmer temperature. The fluctuation of microbial occurrence in a specific region is altered by rainfall although a consistent pattern are not shown (Barata et al., 2012). Yet there is no information on the effect of dry and wet season to imply tropical variations in palm wine composition, which is likely to vary considerably. The variety of regional factors and microbial characteristics determine how a microbial species might adapt to those specific niches. In conjunction to the local climate and specific biogeographical factors, there are limited studies, and further work here will determine the regional typicity of palm wine. It is likely that these factors are important as the interplay among environment, plant species, and winemaking practice impacts the microbial populations and their metabolism and contributes to formation of the final product characteristics, reflecting a fermented beverage product’s origin and terroir (Bokulich et al., 2016; Knight et al., 2015; Liu et al., 2019).

Palm species are a critical aspect that contribute to the differences of palm sap characteristics and thus flavour. There are more than 10 species of palm trees, that have utilized for making palm wine (Hebbar et al., 2018). As showed in **Fig. 2C**, palm species are established and well-adapted to local biogeographical characteristics. For example, *Elaensis* and *Raphia* palm species are used in Africa but unrecorded in both Asia and America. Palm tree specificity can be considered using chemical analysis of the sap where each palm species likely has different composition and quantity of macro and micronutrient, contributing to the growth of microbial communities (Barh & Mazumdar, 2008; Djeni et al., 2022). This variation indicates that palm species genetic diversity and the influence of geographical locality are the significant factors to determine palm wine specificity. These processes and the collated data from our survey are explained in the following sections. Of the studies collated, 17 focussed on description of the microbes present, 7 on the small molecules, which are microbially derived and likely to be sensorially active and 23 studies combined these approaches (**Fig.2B**).

### 3.2 Introduction of microbes into palm sap

Palm sap is believed in a sterile condition upon collection and microbial communities are inoculated through insect visitation and the tapping process (Naknean et al., 2010; Santiago-Urbina et al., 2015; Djeni et al., 2020). Insects and microbes have an intimate relationship with plants, signalling symbiotic associations to help them detecting nutrient, protecting from pathogens, and as parasites (Becher et al., 2018; Douglas, 2011). Insects find sugar source through the detection of released volatile compounds by plants (Becher et al., 2018; Madden et al., 2018). Palm sap is rich in sugars, which attract bees and other insects to the collected sap and can be observed floating in the liquid and visiting the collection vessels. Honeybees seek sugary elements of plants, and then acts as pollinator while vectoring microbes (Evans & Schwarz, 2011). Wasps and vinegar flies are often observed in sugar-rich environments in other fermentations (Lam & Howell, 2015; Stefanini et al., 2012). The close relationships of palm sap and insects reflect the vital role of microbial vectors to contribute to the palm wine flavour characteristics. On the other hands, artisanal processes can also contribute to the introduction of microbial communities. Since the tapping process is carried out with a traditional way, and without deliberate sanitation, the possibility of microbe introduction is inevitable and may be from the collection container, human hands, and cutting equipment such as knives. Microbial loads in palm sap can be influenced by a perpetual tapping practice, which can reduce up to 25% of the sap production (Ziadi et al., 2014). Slicing stress during the tapping process is likely related to the decrease of palm sap nutritive value after 10 weeks consecutive tapping and impacts the microbial load. The decrease of microbial loads directly impacts to the fermentation process, affecting the quality and stability of final product. It is likely that the tapping process is a crucial management step for product hygiene and sustainability of sap production.

## 4 Incidence, interactions, and flavour outcome of palm wine fermentations

The nature of palm wine fermentation is vigorous and complex, where available sugars are rapidly consumed (Das & Tamang, 2023; Djeni et al., 2020). Spontaneous fermentations allow natural microorganisms to deplete organic resources without deliberate addition of selected yeast strains. The fermentation environment is a significant factor, which alters microbial interactions and ecological dynamics (Haloin & Strauss, 2008; Kowallik, 2015). As a result, a change in microbial structure is often observed during the fermentation process. To see microbial diversity and dynamic along the fermentation process, we reanalysed data from published studies conducting next generation sequencing (NGS), which is from 7 abstracted studies with further data from Sumerta et al. (2024); (**Supplementary Table S4**). We subsequently selected data of several studies by inclusion criteria, representing a vast diversity change in 24 hours fermentation (**Fig. 3A & 3B**).

**Fig 3.**
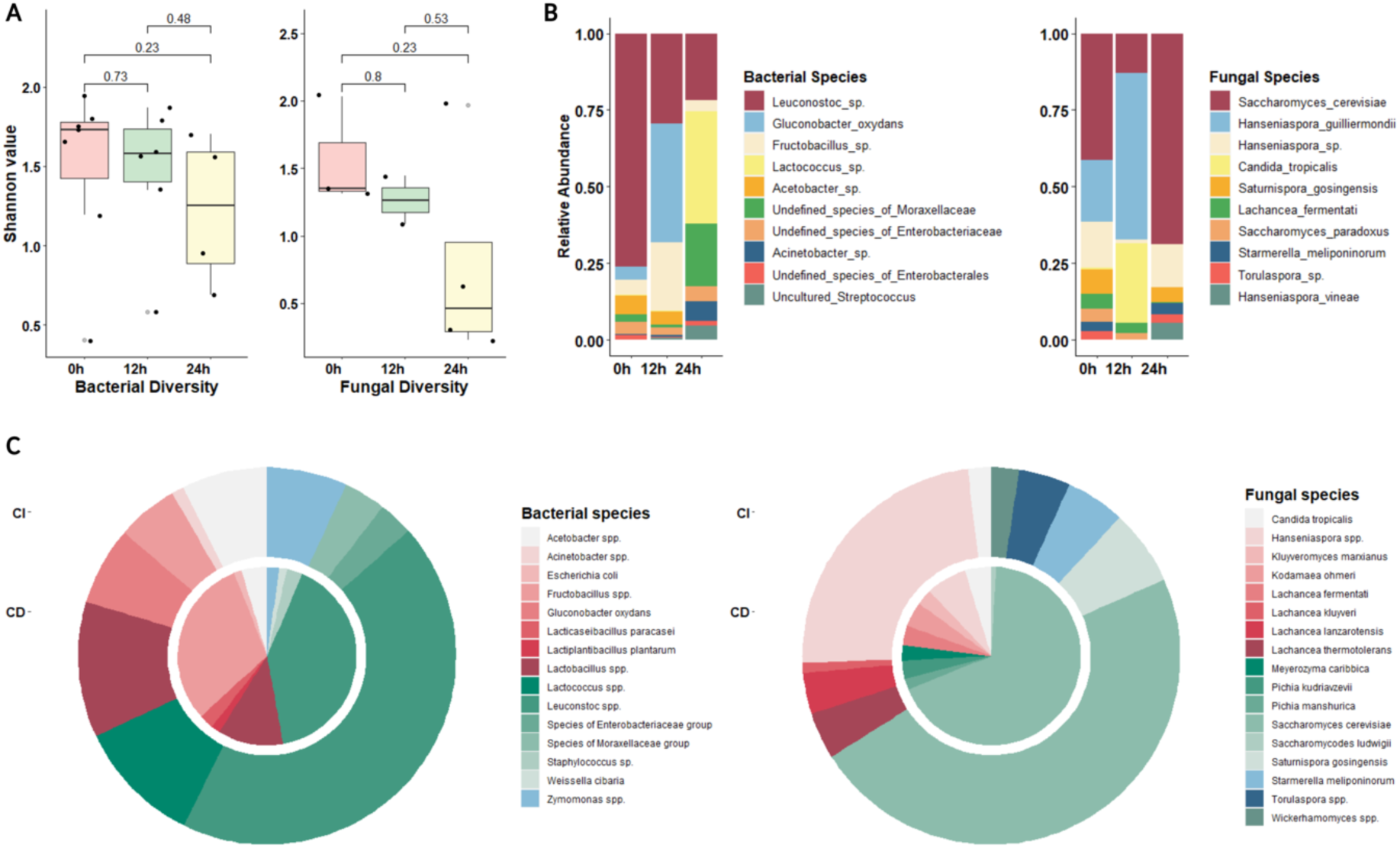
Microbial dynamics and community proportion of palm wine fermentation. Next generation sequencing (NGS) data of 7 abstracted studies with data from Sumerta et al. (2024); **Supplementary Table S4**) were analysed together. A) Microbial diversity loss (measured as Shannon index) over 24 hours palm wine fermentation (Wilcoxon test; p-value <0.05); B) Change in top 10 abundance of microbial species in 24 hours palm wine fermentation from NGS data. C) The proportion of top 10 microbial species, obtained by CD (inner circle; n =33) compared to CI (outer circle; bacteria n=7-1, excluded due to technical limitation; fungi n = 5).

Comparison within country or region, type of sample, and some time frames are less informative because of the limitation in comparable parameters, such as research focus, methodology, and data distribution. Since palm wine fermentation is swift, the species loss of fermenting microbial communities is seen after 24 hours; demonstrated by the decrease in Shannon index during fermentation (**Fig. 3A**; 0, 12, and 24 hours of fermentation). Higher diversity is observed in early stage of the fermentation and decrease to the following process (Alcántara-Hernández et al., 2010). Nwachukwu et al. (2017) used isolation methods and found a diversity loss because of succession of some microbial species. Djeni et al. (2022) reiterated that microbial diversity loss was due to a unique inhabitant species in every stage of palm wine fermentation, that dominates growth.

The relative abundance of bacterial and fungal amplicons from next generation sequencing demonstrates while the initial microbial populations may be maintained for 12 hours, the dominance of *Lactococcus* and *Saccharomyces* is evident (**Fig. 3B**). There are 10 studies that describe a collapse of one microbial group in every stage of palm wine fermentation (**Supplementary Table S6**). Microbial species fluctuate over 24 hours fermentation of palm wine (**Fig. 3B**). The decrease of *Leuconostoc* and the growth of *Lactococcus* present in early stage, with a concurrent drop in *Hanseniaspora* species occurrence and the increase of *Saccharomyces* species. A shift between *Saccharomyces* and non-*Saccharomyces* is inevitable in every stage of the fermentation (Das & Tamang, 2023; Djeni et al., 2022; Gopal et al., 2021; Nwachukwu et al., 2017). This succession of microbial composition over the fermentation process is identified as a key feature to determining palm wine characteristics (Djeni et al., 2022).

### 4.1 *Saccharomyces cerevisiae* is prominent in palm wine fermentations

In a natural fermentation, the complex microbial interactions may result in the dominance of certain species. We sought to find the microbial species most regularly associated with palm fermentation Through the comparison of a conventional microbial isolation and species inferred from amplicon sequencing, we selected data of top 10 abundance of culturable species (CD) (**Supplementary Table S1)** and amplicon sequence variants (CI) (**Supplementary Table S4)**. We compared and found the general pattern of microbial composition in palm wine with the presence of both principal yeast species and principal bacterial genera, which are *S. cerevisiae* and *Leuconostoc*, respectively (**Fig.3C**). Both of methods confirm that the role of *S. cerevisiae* and *Leuconostoc* and this contributes to more than 40% of the total microbial abundance. Some species that do not present in one of those methods may relate to the limitation of each method. Meanwhile, as described in previous studies, *S. cerevisiae* is dominant in all fermentation stages while *Leuconostoc* fluctuates during the final stage (Das & Tamang, 2021; Djeni et al., 2022). The principal species may dominant for the primary product, but non-fermentative or non-*Saccharomyces* species enriches the sensory outputs for fermented beverages more broadly (Li & Liu, 2016; Pietrafesa et al., 2020).

#### 4.1.1 What circumstances allow for the dominance of *S. cerevisiae*?

The role of the key yeast species on producing ethanol as the primary product is pivotal in palm wine fermentation. Some bacterial species are reported to produce ethanol, such as *Zymomonas* spp. (Alcántara-Hernández et al., 2010), but production is less effective than *S. cerevisiae.* The fermentation process is driven by *S. cerevisiae* in the production of many beverages, as this yeast can readily transport sugars, other carbohydrates are hydrolysed, and various compound are produced and released. Some of the produced compounds can inhibit other species and the others yet may display e synergistic actions to adapt with the change of ecological factors during the fermentation process (Ciani et al., 2010; Fleet, 2003). This often leads to the dominance of certain species in a complex spontaneous fermentation of alcoholic beverages of which *S. cerevisiae* in the prime example (Perrone et al., 2013; Pietrafesa et al., 2020; Wang et al., 2015).

*S. cerevisiae* is widely regarded as the most tolerance species to ethanol, acidity, and physical interactions (Ma & Liu, 2010; Henderson & Block, 2014; Albergaria & Arneborg, 2016; Pérez-Torrado et al., 2017). Perrone et al. (2013) suggested that its dominancy is triggered by the occurrence of other species, driving various mechanisms to suppress non-*Saccharomyces*. A regulatory mechanism of its alcohol dehydrogenases (*ADH*) genes controls the ethanol production and at the same time can recover energy when the ethanol excess (Goddard & Greig, 2015). *ADH1* gene regulates the fermentation process to produce ethanol, while *ADH2* gene is responsible to transferring energy from ethanol conversion. Non-*Saccharomyces* species are vulnerable and not competitive in higher alcohol fermentations and are affected by other environmental factors (Pérez-Torrado et al., 2017). *S. cerevisiae* may secrete compounds, such fungicide like mycosin and perform cell-contact abundance and triggering to the death of other species in mixed culture fermentation (Kemsawasd et al., 2015; Wang et al., 2015). Albergaria et al. (2010) found that the 2-10 kDa size of peptides secreted by *S. cerevisiae* can inhibit the growth of *Kluyveromyces marxianus, Kluyveromyces thermotolerans, Torulaspora delbrueckii* and *Hanseniaspora guilliermondii*. The type of cell-to-cell contact and antibiotic production mechanism can also result in the death of certain species (Kemsawasd et al., 2015). Meanwhile, cell-to-cell contact induces the metabolites properties of the species and affect cell viability in late stage of the fermentation (Pietrafesa et al., 2020; Wang et al., 2015). However, physical contact is not the major factor to decrease the presence of non-*Saccharomyces* and confirmed that the metabolites produced by *S. cerevisiae* mostly sharps the non-*Saccharomyces* population (Wang et al., 2015). With the efficiency of energy use, less osmotic stress, high ethanol production, tolerant to lower pH, and enable to inhibit other species, *S. cerevisiae* often survives until the final stage of fermentation (Tra Bi et al., 2019). It is likely that many of these characteristics may also be true of *S. cerevisiae* found in palm wine environments, but these mechanisms have not been directly shown.

#### 4.1.2 Diversity of *Saccharomyces* and the implications for palm wine fermentations

*S. cerevisiae* is a model species for genomics and plays a pivotal role in many food systems, having been selected, adapted, and well domesticated across human history (Aa et al., 2006; Gibbons & Rinker, 2015; Steensels et al., 2019; Steensels & Verstrepen, 2014). Peter et al. (2018) described that the highest variation in *S. cerevisiae* genomes is correlated to the wider differences of phenotypic characters, and it is these characters that are responsible for the signature and originality of fermentation products. The evolutionary process of *S. cerevisiae* at genome level can be modified by human interferences, and the yeasts adapt to various growth conditions, alter phenotypes, and change reproductive mode (Gallone et al., 2016; Liti et al., 2009). In natural conditions, the population structure of *S. cerevisiae* is primarily shaped by ecology (Han et al., 2021). *S. cerevisiae* evolution can be triggered by its response to ecological interactions in harsh fermentation environments (Conacher et al., 2021).

Spontaneous fermentation on fermented beverage making has been reproducibly conducted and if it effects on microbial diversity loss, this potentially leads to a species domestication (Friedrich et al., 2023). Interaction with the natural environment is often possible, where new strains may contribute, and experience variable abiotic conditions (Gallone et al., 2016; Gibbons & Rinker, 2015). Legras et al. (2007) stated that a rich environmental condition, geographical condition, and repetitive use in the same condition results in gradual genotypes evolution, species distinction, and suggest local domestication patterns. However, if spontaneous fermentation of palm wine generates domestication is an open question (Fay & Benavides, 2005). Some reports released that the genome of *S. cerevisiae* populations of African palm wine is admixed compared to global references, showing a unique signature relative to different continents, niches, and favour strain migration through human interference (Ezeronye & Legras, 2009; Tra Bi et al., 2019). Tra Bi et al. (2019) showed that the genetic relationship of *S. cerevisiae* in palm wine is significantly related to ecological niches, that sharing the similar geographical origin generates lower genetic diversity. A local domestication may be attributed to *S. cerevisiae* from palm wine strains, which are observed as homozygous strains (Ezeronye & Legras, 2009; Tapsoba et al., 2015). Further studies with diverse isolates of *S. cerevisiae* genomes will help unravel this question.

### 4.2 Interactions amongst microbes in fermentation

We conducted a network analysis of the co-occurrence of species groups from compiled studies which used culture-dependent methods to assess microbial diversity (**Fig. 4A**). The contribution of *Saccharomyces* is dominant and acts as a central hub to other species group in palm wine fermentation. It also has a strong relationship to other yeast genera (*Hanseniaspora, Candida*, and *Pichia*) and the core bacteria in organic acids production, such as *Lactobacillus*, *Leuconostoc*, and acetic acid bacteria (AAB). These show a vital connection pattern of those microbial groups on constructing palm wine characters and suggest a fermentation model retrieved from multi-geographical studies. This pattern reveals a network of microbial ecology, and reveals potential microbial interactions which may be able to predict the occurrence of unwanted species (Fuhrman, 2009).

**Fig 4.**
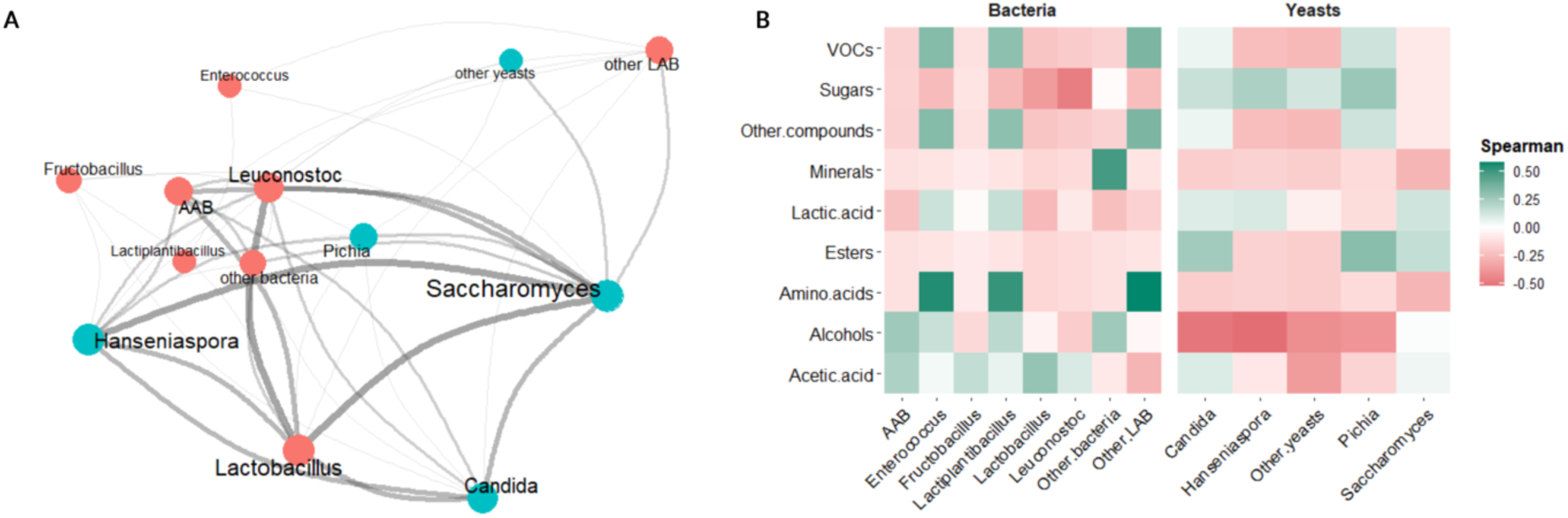
Co-occurrence pattern of microbe-microbe and microbe-metabolite relationships on palm wine fermentation. A) Network analysis of the relationship of microbial group in palm wine fermentation. The contribution of each group is represented as an edge. Different colour showing different microbial group (blue= yeasts, red= bacteria), then different size of font represents the abundance value. The data set used for this study consists of 23 studies based on the abundance and frequency each species group in **Supplementary Table S3**, omitting 7 studies of NGS, where data was abstracted in **Supplementary Table S5** and used R studio adopting the codes by Schweinberger (2024). B) Correlation heatmap between microbial and metabolite groups (n= 23-3, studies which used amplicon sequencing were disregarded). The frequency of microbial and metabolite groups was abstracted from **Supplementary Table S3** and analysed with Spearman correlations (p value > 0.05).

In spontaneous fermentation, interactions have a significant role on driving the fermentation process, involving various microbial groups. Interactions among microbial species may deliver positive and negative impacts, which determine the product formation (Ciani & Comitini, 2015; Cordero & Datta, 2016; Viljoen, 2001; Zuñiga et al., 2017). In spontaneous fermentations, synergistic interactions among different species mediate tolerance to the robustness of dynamic condition in particular level. Mutualism and commensalism interaction are attributed to the synergistic interaction with cross-feeding mechanism or supporting certain metabolite producer (Silverstein et al., 2023; Smid & Lacroix, 2013). For instance, yeasts ferment the sugar into ethanol while acetic acid bacteria convert the ethanol into acetic acid which is not affect the yeast (De Vuyst & Leroy, 2020). Lactic acid bacteria and yeast sometimes support each other by exchanging some metabolites for their metabolism (Smid & Lacroix, 2013). Each group of microbial species has different way to develop its functionality by interacting with other species. Fleet (2003) highlighted the interaction among yeast species contributes to the fermentation behaviour while yeast-bacteria interactions determine the ecological dynamic. This interesting feature of complex microbial communities interaction can be proposed as new model of the fermentation to create flavourful product (Ciani et al., 2010). Specific interactions between microbes have not been described in a palm wine environment, but they are likely to occur.

### 4.3 The implications of metabolite release on fermentation dynamics

The associations of microbial communities and chemical release have a major contribution on constructing palm wine characteristics. A wide range of microbial groups occupies in every stage of the fermentation process but following the depletion of sugar in the fermentation process, microbial diversity decreases because of biochemicals alteration and ecological interactions. Here, we abstracted 10 studies (**Supplementary Table S6**), exploring both in microbial communities and chemical composition along the fermentation stages, representing palm wine fermentation dynamics and summarised in **Fig. 5**. The abundance of ethanol is likely attributed to the presence of *Saccharomyces* yeasts during fermentation. Lactic acid bacteria (LAB) and acetic acid bacteria (AAB) grow on palm wine fermentation related to the significant number of organic acids in final stages of fermentation (Adamu-Governor et al., 2018; Kouame et al., 2020; Ouoba et al., 2012; Somashekaraiah et al., 2019). Total phenolics and flavonoids impact shifts in microbial community composition through the fermentation process (Das & Tamang, 2023). We presented the examples of chemical yield in the table, which represents the gradual change in line graph. It is important to note that chemical content of palm wine ultimately depends on, geographical location, palm species, and the way analytical process and research design conducted, so the results vary among studies. By summarising related studies, the general pattern was demonstrated to provide a better understanding of palm wine fermentation.

**Fig. 5.**
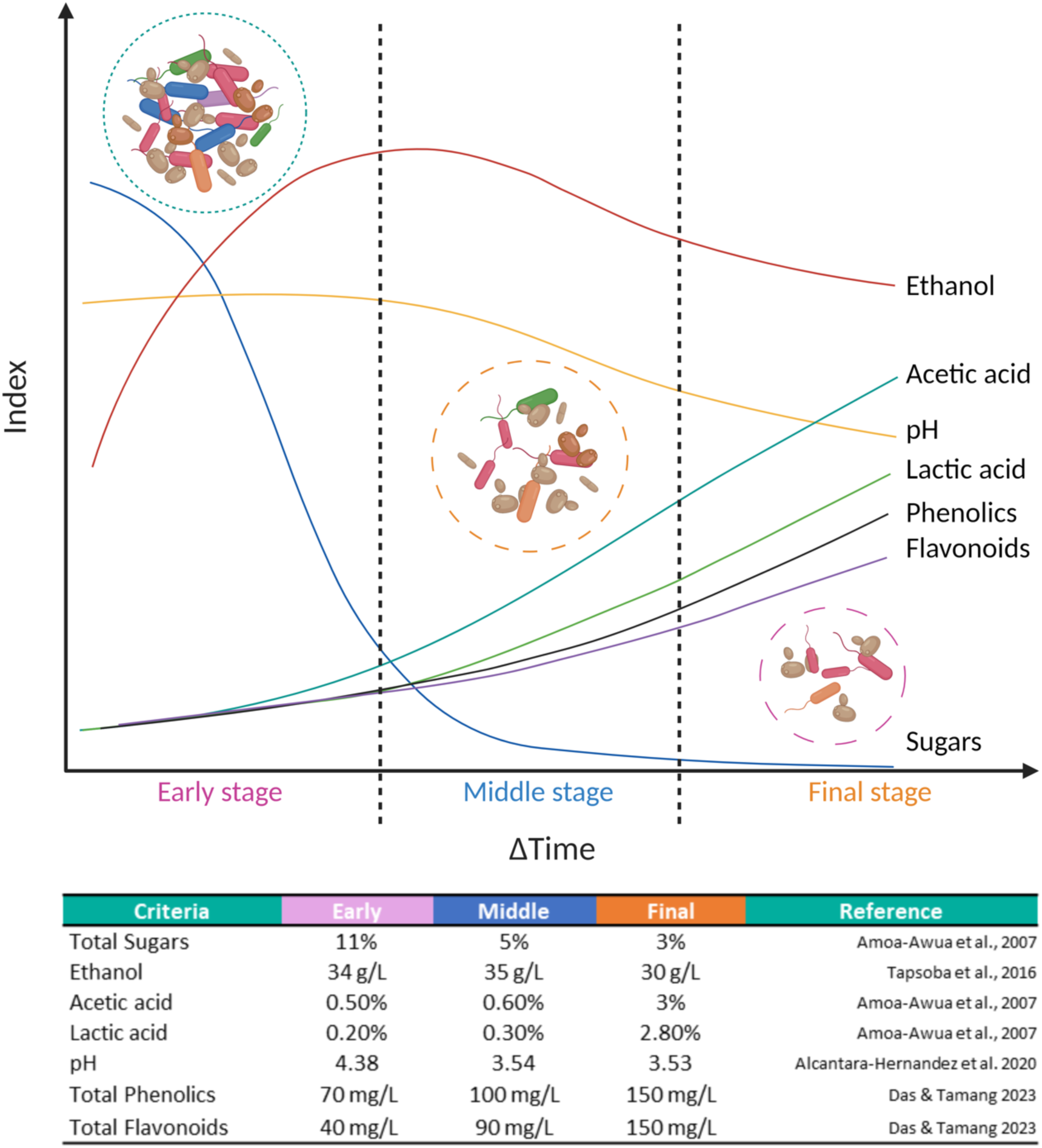
Schematic of the process of palm wine fermentation which summarises the microbial and chemical composition, dynamics of microbial communities, raw materials, and metabolites over time. A gradual pattern of palm wine fermentation was reconstructed from compiled studies (n= 10, **Supplementary Table S6**) and table shows the examples of chemical yield during the fermentation process, created with BioRender.com.

We simulated the co-occurrence of microbial groups and compounds based on the correlation analysis of each microbial group follows the biochemical pattern (**Fig. 4B**). Data was summarised from the frequency of microbial groups and metabolite components presented in each study. Lactic and acetic acid production in the final stage of palm wine fermentation positively corelate to the frequency of lactic acid and acetic acid bacteria group while *Saccharomyces* and *Candida* group may also have similar contribution. These compounds reduce the pH and alter the microbial diversity. Since pH decreasing, the evaporation of volatile fatty acids is possibly high (Derikx et al., 1994). Meanwhile in alcohol production, non-*Saccharomyces* yeasts have a negative correlation compared to bacterial groups and *Saccharomyces*. This simulation provides the influence of metabolite components on the composition of microbial diversity, which can demonstrate the fermentation dynamics.

## 5 The intersection of culture for palm wine production in modern food systems

The process of palm wine making is still performed in traditional ways from tapping process to the fermentation. Depend on palm species, tapping process can be conducted through destructive and non-destructive methods (Djeni et al., 2020). In West Africa, destructive tapping process is mostly observed by notching or perforating the apical meristem, often resulting in the death of the tree. The widely used method is non-destructive by cutting the inflorescence and the sap drips from cutting tip to the collection container (Limtong et al., 2020). The collected palm sap is then processed into various products for daily consumption or sold. For some areas, this activity is the main job of local communities and maintains economic stability in many villages.

The use of palm sap and/or wine has an important role as part of culture and tradition (Wijaya et al., 2024). Palm spirits can be used as part of wedding ceremonies, event celebration, welcoming rituals, and religious rituals. In some tropical countries, palm sap products are also responsible for ethnomedicines. Anthropologists have acknowledged that palm wine is important as stimulant, depressant, and part of traditional medication, well-written in traditional transcripts (Jakl, 2021). Recent scientific reports also suggested that palm sap/wine compounds have health benefits, related to the empirical thoughts of local people, such as decrease diabetes risk, antioxidant, antibiotics, anaemia treatment, and immunomodulators (Barh & Mazumdar, 2008; Hebbar et al., 2018; Laklaeng & Kwanhian, 2020; Leida et al., 2020; Mia et al., 2020; Senghoi & Klangbud, 2021). Palm wine contains phenolic compounds that may contribute to antioxidant activity in the plasma (Hebbar et al., 2018; Xia et al., 2011) and modulate the growth of gut microbiota (Cardona et al., 2013; Nash et al., 2018; Queipo-Ortuño et al., 2012). A probiotic-type microbiome that benefits for digestive system is also found in fresh palm sap (Gopal et al., 2021) and fermented sap showed bioactive chemical activities (Kurniawan et al., 2018). We suggest that basic information on microbial community structure and dynamic of metabolite profile benefits local and traditional practices, that support regional identity and preservation of traditional practices.

Understanding microbial ecology and controlling microbial composition are pivotal to beverage quality and a major challenge for future product development (Gobert et al., 2022). In general, inconsistent sensory characteristics results from spontaneous fermentation as it is difficult to control product quality and consistency. The presence of spoilage bacteria is inevitable amid a spontaneous fermentation, for example lactic acid and acetic acid bacteria, that can compete for nutrients and produce lactic and acetic acids, which limit the ethanol production during the fermentation process (Beckner et al., 2011). Intervention at harvesting may obtain higher ethanol production before the spoilage bacteria rapidly conducts the acidic fermentation. Deploying defined starter cultures may obtain consistent, efficient, and desired quality of the alcohol products; however, the end-products might lack complexity of organoleptic characteristics (Steensels & Verstrepen, 2014). The fermentation process has evolved to current commercial practices with selected yeast strains (Cavalieri et al., 2003; Steensels & Verstrepen, 2014) to enhance aroma and flavour (Medina et al., 2013; Rodríguez-Lerma et al., 2011). Although this practice yet applicable in palm wine, co-culturing with selected strain could be considered in the future. We recognise there is a need to retain and foster traditional food products, which may at odds with the use of commercial starter cultures.

## 6 Conclusions, limitations, and prospects

Palm wine is a unique and culturally important beverage. The fermentation process is challenged by many factors, which contribute to differences of product composition and distinction. Technologically behind from the production of grape and rice wines, there is no standardized procedure widely used to produce palm wine, which remains at extremely small-scale and conducted with traditional methods. We have defined the facets of palm wine fermentation and composition reflecting the importance of ecological specificity on beverage characteristics but encourage addition of further studies across other regions and ecological variations to add to the knowledge of this unique beverage. Further studies will help typical geographical features and help preserve traditional knowledge and distinctive foods for benefit of local communities. We support further actions to strengthen fundamental understandings of palm wine production to support sustainable product development and appreciating local knowledges and traditions.

## Acknowledgements

Authors would like to pass our gratitude to University of Melbourne for access to the high-performance computing for faster analysis. Theodore N. Djeni and Kumaraswamy Jeyaram who provided the sequencing data available publicly for compiled analysis. SSEAC (Sydney Southeast Asia Centre) for the intensive writing program, supporting on the writing process.

## Author contributions

INS: research design, data compilation, analysis, writing and editing the manuscript

XR: data analysis supervision and review the manuscript

KH: research design supervision, writing, review and editing the manuscript

## References

Aa, E., Townsend, J. P., Adams, R. I., Nielsen, K. M., & Taylor, J. W. (2006). Population structure and gene evolution in Saccharomyces cerevisiae. FEMS Yeast Research, 6(5), 702–715. 10.1111/j.1567-1364.2006.00059.x

Adamu-Governor, O. L., Shittu, T. A., Afolabi, O. R., & Uzochukwu, S. V. A. (2018). Screening for gum-producing Lactic acid bacteria in Oil palm (Elaeis guineensis) and raphia palm (Raphia regalis) sap from South-West Nigeria. Food Science & Nutrition, 6(8), 2047–2055. 10.1002/fsn3.750

Albergaria, H., & Arneborg, N. (2016). Dominance of Saccharomyces cerevisiae in alcoholic fermentation processes: Role of physiological fitness and microbial interactions. Applied Microbiology and Biotechnology, 100(5), 2035–2046. 10.1007/s00253-015-7255-0

Albergaria, H., Francisco, D., Gori, K., Arneborg, N., & Gírio, F. (2010). Saccharomyces cerevisiae CCMI 885 secretes peptides that inhibit the growth of some non-Saccharomyces wine-related strains. Applied Microbiology and Biotechnology, 86(3), 965–972. 10.1007/s00253-009-2409-6

Alcántara-Hernández, R. J., Rodríguez-Álvarez, J. A., Valenzuela-Encinas, C., Gutiérrez-Miceli, F. A., Castañón-González, H., Marsch, R., Ayora-Talavera, T., & Dendooven, L. (2010). The bacterial community in ‘taberna’ a traditional beverage of Southern Mexico. Letters in Applied Microbiology, 51(5), 558–563. 10.1111/j.1472-765X.2010.02934.x

Amoa-Awua, W. K., Sampson, E., & Tano-Debrah, K. (2007). Growth of yeasts, lactic and acetic acid bacteria in palm wine during tapping and fermentation from felled oil palm (Elaeis guineensis) in Ghana. Journal of Applied Microbiology, 102(2), 599–606. 10.1111/j.1365-2672.2006.03074.x

Anderson, K., & Wittwer, G. (2015). Asia’s evolving role in global wine markets. China Economic Review, 35, 1–14. 10.1016/j.chieco.2015.05.003

Astudillo-Melgar, F., Ochoa-Leyva, A., Utrilla, J., & Huerta-Beristain, G. (2019). Bacterial Diversity and Population Dynamics During the Fermentation of Palm Wine From Guerrero Mexico. Frontiers in Microbiology, 10, 531. 10.3389/fmicb.2019.00531

Barata, A., Malfeito-Ferreira, M., & Loureiro, V. (2012). The microbial ecology of wine grape berries. International Journal of Food Microbiology, 153(3), 243–259. 10.1016/j.ijfoodmicro.2011.11.025

Barh, D., & Mazumdar, B. C. (2008). Comparative Nutritive Values of Palm Saps Before and after Their Partial Fermentation and Effective Use of Wild Date (Phoenix sylvestris Roxb.) Sap in Treatment of Anemia. Med. Sci., 3(2), 173–176.

Becher, P. G., Hagman, A., Verschut, V., Chakraborty, A., Rozpędowska, E., Lebreton, S., Bengtsson, M., Flick, G., Witzgall, P., & Piškur, J. (2018). Chemical signaling and insect attraction is a conserved trait in yeasts. Ecology and Evolution, 8(5), 2962–2974. 10.1002/ece3.3905

Beckner, M., Ivey, M. L., & Phister, T. G. (2011). Microbial contamination of fuel ethanol fermentations: Bioethanol contamination. Letters in Applied Microbiology, 53(4), 387–394. 10.1111/j.1472-765X.2011.03124.x

Bokulich, N. A., Collins, T. S., Masarweh, C., Allen, G., Heymann, H., Ebeler, S. E., & Mills, D. A. (2016). Associations among Wine Grape Microbiome, Metabolome, and Fermentation Behavior Suggest Microbial Contribution to Regional Wine Characteristics. mBio, 7(3), e00631–16. 10.1128/mBio.00631-16

Candiago, S., Tscholl, S., Bassani, L., Fraga, H., & Egarter Vigl, L. (2022). A geospatial inventory of regulatory information for wine protected designations of origin in Europe. Scientific Data, 9(1), Article 1. 10.1038/s41597-022-01513-0

Cardona, F., Andrés-Lacueva, C., Tulipani, S., Tinahones, F. J., & Queipo-Ortuño, M. I. (2013). Benefits of polyphenols on gut microbiota and implications in human health. The Journal of Nutritional Biochemistry, 24(8), 1415–1422. 10.1016/j.jnutbio.2013.05.001

Cavalieri, D., McGovern, P. E., Hartl, D. L., Mortimer, R., & Polsinelli, M. (2003). Evidence for S. cerevisiae fermentation in ancient wine. Journal of Molecular Evolution, 57 *Suppl 1*, S226–232. 10.1007/s00239-003-0031-2

Chandrasekhar, K., Sreevani, S., Seshapani, P., & Pramodhakumari, J. (2012). A Review on palm wine. 2(1), 33–38.

Ciani, M., & Comitini, F. (2015). Yeast interactions in multi-starter wine fermentation. Current Opinion in Food Science, 1, 1–6. 10.1016/j.cofs.2014.07.001

Ciani, M., Comitini, F., Mannazzu, I., & Domizio, P. (2010). Controlled mixed culture fermentation: A new perspective on the use of non-Saccharomyces yeasts in winemaking. FEMS Yeast Research, 10(2), 123–133. 10.1111/j.1567-1364.2009.00579.x

Ciani, M., Stringini, M., & Comitini, F. (2012). Palm Wine. In Handbook of Plant-Based Fermented Food and Beverage Technology (pp. 652–659). CRC Press. 10.1201/b12055-41

Conacher, C., Luyt, N., Naidoo-Blassoples, R., Rossouw, D., Setati, M., & Bauer, F. (2021). The ecology of wine fermentation: A model for the study of complex microbial ecosystems. Applied Microbiology and Biotechnology, 105(8), 3027–3043. 10.1007/s00253-021-11270-6

Corbin, K. R., Hsieh, Y. S. Y., Betts, N. S., Byrt, C. S., Henderson, M., Stork, J., DeBolt, S., Fincher, G. B., & Burton, R. A. (2015). Grape marc as a source of carbohydrates for bioethanol: Chemical composition, pre-treatment and saccharification. Bioresource Technology, 193, 76–83. 10.1016/j.biortech.2015.06.030

Cordero, O. X., & Datta, M. S. (2016). Microbial interactions and community assembly at microscales. Current Opinion in Microbiology, 31, 227–234. 10.1016/j.mib.2016.03.015

Coutiño, B., Flores, A. C., Vela-Gutiérrez, G., Sepúlveda, L., Aguilar, C. N., Chavez-Gonzalez, M., & Rodríguez, R. (2020). 7 - Tavern or Coyol Wine: A Beverage From Palm Sap With Biotechnological Potential. In A. M. Grumezescu & A. M. Holban (Eds.), Biotechnological Progress and Beverage Consumption (pp. 233–252). Academic Press. 10.1016/B978-0-12-816678-9.00007-2

Das, S., & Tamang, J. P. (2021). Changes in microbial communities and their predictive functionalities during fermentation of toddy, an alcoholic beverage of India. Microbiological Research, 248, 126769. 10.1016/j.micres.2021.126769

Das, S., & Tamang, J. P. (2023). Fermentation Dynamics of Naturally Fermented Palm Beverages of West Bengal and Jharkhand in India. Fermentation, 9(3), Article 3. 10.3390/fermentation9030301

De Guidi, I., Legras, J.-L., Galeote, V., & Sicard, D. (2023). Yeast domestication in fermented food and beverages: Past research and new avenues. Current Opinion in Food Science, 101032. 10.1016/j.cofs.2023.101032

Derikx, P. J. L., Willers, H. C., & ten Have, P. J. W. (1994). Effect of pH on the behaviour of volatile compounds in organic manures during dry-matter determination. Bioresource Technology, 49(1), 41–45. 10.1016/0960-8524(94)90171-6

Deselnicu, O. C., Costanigro, M., Souza-Monteiro, D. M., & McFadden, D. T. (2013). A Meta-Analysis of Geographical Indication Food Valuation Studies: What Drives the Premium for Origin-Based Labels? Journal of Agricultural and Resource Economics, 38(2), 204–219.

De Vuyst, L., & Leroy, F. (2020). Functional role of yeasts, lactic acid bacteria and acetic acid bacteria in cocoa fermentation processes. FEMS Microbiology Reviews, 44(4), 432–453. 10.1093/femsre/fuaa014

Djeni, T. N., Keisam, S., Kouame, K. H., Assohoun-Djeni, C. N., Ake, F. D. M., Amoikon, L. S. T., Tuikhar, N., Labala, R. K., Dje, M. K., & Jeyaram, K. (2022). Dynamics of microbial populations and metabolites of fermenting saps throughout tapping process of ron and oil palm trees in Côte d’Ivoire. Frontiers in Microbiology, 13. https://www.frontiersin.org/articles/10.3389/fmicb.2022.954917

Djeni, T. N., Kouame, K. H., Ake, F. D. M., Amoikon, L. S. T., Dje, M. K., & Jeyaram, K. (2020). Microbial Diversity and Metabolite Profiles of Palm Wine Produced From Three Different Palm Tree Species in Côte d’Ivoire. Scientific Reports, 10(1), Article 1. 10.1038/s41598-020-58587-2

Doddipalla, R., Rendedula, D., Ganneru, S., Kaliyaperumal, M., & Mudiam, M. K. R. (2022). Understanding metabolic perturbations in palm wine during storage using multi-platform metabolomics. LWT, 155, 112889. 10.1016/j.lwt.2021.112889

Douglas, A. E. (2011). Lessons from Studying Insect Symbioses. Cell Host & Microbe, 10(4), 359–367. 10.1016/j.chom.2011.09.001

Evans, J. D., & Schwarz, R. S. (2011). Bees brought to their knees: Microbes affecting honey bee health. Trends in Microbiology, 19(12), 614–620. 10.1016/j.tim.2011.09.003

Ezeronye, O. u., & Legras, J.-L. (2009). Genetic analysis of Saccharomyces cerevisiae strains isolated from palm wine in eastern Nigeria. Comparison with other African strains. Journal of Applied Microbiology, 106(5), 1569–1578. 10.1111/j.1365-2672.2008.04118.x

Fay, J. C., & Benavides, J. A. (2005). Evidence for domesticated and wild populations of Saccharomyces cerevisiae. PLoS Genetics, 1(1), 66–71. 10.1371/journal.pgen.0010005

Fleet, G. H. (2003). Yeast interactions and wine flavour. International Journal of Food Microbiology, 86(1), 11–22. 10.1016/S0168-1605(03)00245-9

Fontoin, H., Saucier, C., Teissedre, P.-L., & Glories, Y. (2008). Effect of pH, ethanol and acidity on astringency and bitterness of grape seed tannin oligomers in model wine solution. Food Quality and Preference, 19(3), 286–291. 10.1016/j.foodqual.2007.08.004

Friedrich, A., Gounot, J.-S., Tsouris, A., Bleykasten, C., Freel, K., Caradec, C., & Schacherer, J. (2023). Contrasting genomic evolution between domesticated and wild Kluyveromyces lactis yeast populations. Genome Biology and Evolution, 15(2), evad004. 10.1093/gbe/evad004

Fuhrman, J. A. (2009). Microbial community structure and its functional implications. Nature, 459(7244), Article 7244. 10.1038/nature08058

Gallone, B., Steensels, J., Prahl, T., Soriaga, L., Saels, V., Herrera-Malaver, B., Merlevede, A., Roncoroni, M., Voordeckers, K., Miraglia, L., Teiling, C., Steffy, B., Taylor, M., Schwartz, A., Richardson, T., White, C., Baele, G., Maere, S., & Verstrepen, K. J. (2016). Domestication and Divergence of Saccharomyces cerevisiae Beer Yeasts. Cell, 166(6), 1397–1410.e16. 10.1016/j.cell.2016.08.020

Gibbons, J. G., & Rinker, D. C. (2015). The genomics of microbial domestication in the fermented food environment. Current Opinion in Genetics & Development, 35, 1–8. 10.1016/j.gde.2015.07.003

Gobert, A., Evers, M. S., Morge, C., Sparrow, C., & Delafont, V. (2022). Comparison of DNA purification methods for high-throughput sequencing of fungal communities from wine fermentation. MicrobiologyOpen, 11(5), e1321. 10.1002/mbo3.1321

Goddard, M. R., & Greig, D. (2015). Saccharomyces cerevisiae: A nomadic yeast with no niche? FEMS Yeast Research, 15(3). 10.1093/femsyr/fov009

González, Á., & De Vuyst, L. (2009). Vinegars from Tropical Africa. In L. Solieri & P. Giudici (Eds.), Vinegars of the World (pp. 209–221). Springer Milan. 10.1007/978-88-470-0866-3_13

Gopal, M., Shil, S., Gupta, A., Hebbar, K. B., & Arivalagan, M. (2021). Metagenomic Investigation Uncovers Presence of Probiotic-Type Microbiome in Kalparasa® (Fresh Unfermented Coconut Inflorescence Sap). Frontiers in Microbiology, 12, 662783. 10.3389/fmicb.2021.662783

Gray, N. D., & Head, I. M. (2008). Microbial Ecology. In S. E. Jørgensen & B. D. Fath (Eds.), Encyclopedia of Ecology (pp. 2357–2368). Academic Press. 10.1016/B978-008045405-4.00519-X

Gregorio, C. G. C. (2023). Philippine Traditional Alcoholic Beverages: A Germinal Study. In J.-M. Mérillon, C. Riviere, & G. Lefèvre (Eds.), Natural Products in Beverages (pp. 1–30). Springer International Publishing. 10.1007/978-3-031-04195-2_188-1

Haloin, J. R., & Strauss, S. Y. (2008). Interplay between ecological communities and evolution: Review of feedbacks from microevolutionary to macroevolutionary scales. Annals of the New York Academy of Sciences, 1133, 87–125. 10.1196/annals.1438.003

Han, D.-Y., Han, P.-J., Rumbold, K., Koricha, A. D., Duan, S.-F., Song, L., Shi, J.-Y., Li, K., Wang, Q.-M., & Bai, F.-Y. (2021). Adaptive Gene Content and Allele Distribution Variations in the Wild and Domesticated Populations of Saccharomyces cerevisiae. Frontiers in Microbiology, 12, 631250. 10.3389/fmicb.2021.631250

Hebbar, K. B., Pandiselvam, R., Manikantan, M. R., Arivalagan, M., Beegum, S., & Chowdappa, P. (2018). Palm Sap—Quality Profiles, Fermentation Chemistry, and Preservation Methods. Sugar Tech, 20(6), 621–634. 10.1007/s12355-018-0597-z

Henderson, C. M., & Block, D. E. (2014). Examining the Role of Membrane Lipid Composition in Determining the Ethanol Tolerance of Saccharomyces cerevisiae. Applied and Environmental Microbiology, 80(10), 2966–2972. 10.1128/AEM.04151-13

Hermansyah, H., Novia, N., Minetaka, S., & Satoshi, H. (2015). Candida tropicalis Isolated from Tuak, a North Sumatera-Indonesian Traditional Beverage, for Bioethanol Production. Microbiology and Biotechnology Letters, 43(3), 241–248. 10.4014/mbl.1506.06002

Ikeh, I. M., Anele, B. C., Ukanwa, C. C., & Njoku, S. O. (2021). Analysis of the Microbial Quality of Locally Consumed Palm Wine Sold in Elele Community of Rivers State Nigeria. European Journal of Nutrition & Food Safety, 62–69. 10.9734/ejnfs/2021/v13i730437

Ivanova, V., Stefova, M., & Chinnici, F. (2010). Determination of the polyphenol contents in Macedonian grapes and wines by standardized spectrophotometric methods. Journal of the Serbian Chemical Society, 75(1), 45–59. 10.2298/JSC1001045I

Jakl, J. (2021). Alcohol in early Java: Its social and cultural significance. Brill.

Jespersen, L. (2003). Occurrence and taxonomic characteristics of strains of Saccharomyces cerevisiae predominant in African indigenous fermented foods and beverages. Fems Yeast Research, 3(2), 191–200. 10.1016/S1567-1356(02)00185-X

Karamoko, D., Djeni, N. T., N’guessan, K. F., Bouatenin, K. M. J.-P., & Dje, K. M. (2012). The biochemical and microbiological quality of palm wine samples produced at different periods during tapping and changes which occured during their storage. Food Control, 26(2), 504–511. 10.1016/j.foodcont.2012.02.018

Kemsawasd, V., Branco, P., Almeida, M. G., Caldeira, J., Albergaria, H., & Arneborg, N. (2015). Cell-to-cell contact and antimicrobial peptides play a combined role in the death of Lachanchea thermotolerans during mixed-culture alcoholic fermentation with Saccharomyces cerevisiae. FEMS Microbiology Letters, 362. 10.1093/femsle/fnv103

Knight, S., Klaere, S., Fedrizzi, B., & Goddard, M. R. (2015). Regional microbial signatures positively correlate with differential wine phenotypes: Evidence for a microbial aspect to terroir. Scientific Reports, 5(1), Article 1. 10.1038/srep14233

Kouame, H. K., Ake, M. D. F., Assohoun, N. M. C., Dje, M. K., & Djeni, N. T. (2020). Dynamics and species diversity of lactic acid bacteria involved in the spontaneous fermentation of various palm tree saps during palm wine tapping in Cote d’Ivoire. World Journal of Microbiology & Biotechnology, 36(5), 64. 10.1007/s11274-020-02832-3

Kowallik, V. (2015). The natural ecology of Saccharomyces yeasts [PhD Thesis]. the Christian Albrechts University of Kiel.

Kurniawan, T., Jayanudin, J., Kustiningsih, I., & Adha Firdaus, M. (2018). Palm Sap Sources, Characteristics, and Utilization in Indonesia. Journal of Food and Nutrition Research, 6(9), 590–596. 10.12691/jfnr-6-9-8

Laklaeng, S., & Kwanhian, W. (2020). Immunomodulation Effect of Nypa fruticans Palm Vinegar. Walailak Journal of Science and Technology (WJST), 17(11), 1200–1210. 10.48048/wjst.2020.10719

Lam, S. S. T. H., & Howell, K. S. (2015). Drosophila-associated yeast species in vineyard ecosystems. FEMS Microbiology Letters, 362(20), fnv170. 10.1093/femsle/fnv170

Lee, T. J., Liu, Y., Liu, W.-A., Lin, Y.-F., Lee, H.-H., Ke, H.-M., Huang, J.-P., Lu, M.-Y. J., Hsieh, C.-L., Chung, K.-F., Liti, G., & Tsai, I. J. (2022). Extensive sampling of Saccharomyces cerevisiae in Taiwan reveals ecology and evolution of predomesticated lineages. Genome Research, gr.276286.121. 10.1101/gr.276286.121

Legras, J.-L., Merdinoglu, D., Cornuet, J.-M., & Karst, F. (2007). Bread, beer and wine: Saccharomyces cerevisiae diversity reflects human history. Molecular Ecology, 16(10), 2091–2102. 10.1111/j.1365-294X.2007.03266.x

Leida, I. M., Thaha, R. M., Yusnitasari, A. S., & Afsahyana. (2020). Effect of Sap Palm (Borassus flabellifer) on Blood Glucose Level in Pre-Diabetic Patients. International Journal of Current Research and Review, 12(24), 96–100. 10.31782/IJCRR.2020.122419

Li, X., & Liu, S.-Q. (2016). Antagonism between *Saccharomyces cerevisiae* and *Williopsis mrakii* and significance for mixed yeast alcoholic beverage fermentations. International Journal of Food Science & Technology, 51(3), 656–663. 10.1111/ijfs.13025

Limtong, S., Am-In, S., Kaewwichian, R., Kaewkrajay, C., & Jindamorakot, S. (2020). Exploration of yeast communities in fresh coconut, palmyra, and nipa palm saps and ethanol-fermenting ability of isolated yeasts. Antonie van Leeuwenhoek, 113(12), 2077–2095. 10.1007/s10482-020-01479-2

Liti, G., Carter, D. M., Moses, A. M., Warringer, J., Parts, L., James, S. A., Davey, R. P., Roberts, I. N., Burt, A., Koufopanou, V., Tsai, I. J., Bergman, C. M., Bensasson, D., O’Kelly, M. J. T., van Oudenaarden, A., Barton, D. B. H., Bailes, E., Nguyen, A. N., Jones, M., … Louis, E. J. (2009). Population genomics of domestic and wild yeasts. Nature, 458(7236), 337–341. 10.1038/nature07743

Liu, D., Chen, Q., Zhang, P., Chen, D., & Howell, K. S. (2020). The Fungal Microbiome Is an Important Component of Vineyard Ecosystems and Correlates with Regional Distinctiveness of Wine. mSphere, 5(4), e00534–20. 10.1128/mSphere.00534-20

Liu, D., Zhang, P., Chen, D., & Howell, K. (2019). From the Vineyard to the Winery: How Microbial Ecology Drives Regional Distinctiveness of Wine. Frontiers in Microbiology, 10, 2679. 10.3389/fmicb.2019.02679

Ma, M., & Liu, Z. L. (2010). Mechanisms of ethanol tolerance in Saccharomyces cerevisiae. Applied Microbiology and Biotechnology, 87(3), 829–845. 10.1007/s00253-010-2594-3

Madden, A. A., Epps, M. J., Fukami, T., Irwin, R. E., Sheppard, J., Sorger, D. M., & Dunn, R. R. (2018). The ecology of insect–yeast relationships and its relevance to human industry. Proceedings of the Royal Society B: Biological Sciences, 285(1875), 20172733. 10.1098/rspb.2017.2733

Malek, G., Moktar, H., Luciano, B., Vittorio, C., & Salvatore, M. (2017). Identification of acetic acid bacteria isolated from Tunisian palm sap. African Journal of Microbiology Research, 11(15), 596–602. 10.5897/AJMR2016.8247

Medina, K., Boido, E., Fariña, L., Gioia, O., Gomez, M. E., Barquet, M., Gaggero, C., Dellacassa, E., & Carrau, F. (2013). Increased flavour diversity of Chardonnay wines by spontaneous fermentation and co-fermentation with Hanseniaspora vineae. Food Chemistry, 141(3), 2513–2521. 10.1016/j.foodchem.2013.04.056

Mia, M. A.-T., Mosaib, M. G., Khalil, M. I., Islam, M. A., & Gan, S. H. (2020). Potentials and Safety of Date Palm Fruit against Diabetes: A Critical Review. Foods, 9(11), 1557. 10.3390/foods9111557

Moore, H. E., & Uhl, N. W. (2019, March 19). Palm. Palm. https://www.britannica.com/plant/palm-tree

Naknean, P., Meenune, M., & Roudaut, G. (2010). Characterization of palm sap harvested in Songkhla province, Southern Thailand. International Food Research Journal, 17, 977–986.

Nash, V., Ranadheera, C. S., Georgousopoulou, E. N., Mellor, D. D., Panagiotakos, D. B., McKune, A. J., Kellett, J., & Naumovski, N. (2018). The effects of grape and red wine polyphenols on gut microbiota – A systematic review. Food Research International, 113, 277–287. 10.1016/j.foodres.2018.07.019

Nwachukwu, I., Ibekwe, V. I., & Anyanwu, B. N. (2017). Investigation of physicochemical and microbial succession parameters of palm wine. 4(4).

Nwaiwu, O., Ibekwe, V. I., Amadi, E. S., Udebuani, A. C., Nwanebu, F. C., Oguoma, O. I., & Nnokwe, J. C. (2016). Evaluation of Fermentation Products of Palm Wine Yeasts and Role of Sacoglottis gabonensis Supplement on Products Abundance. Beverages, 2(2), Article 2. 10.3390/beverages2020009

Ouoba, L. i. i., Kando, C., Parkouda, C., Sawadogo-Lingani, H., Diawara, B., & Sutherland, J. p. (2012). The microbiology of Bandji, palm wine of Borassus akeassii from Burkina Faso: Identification and genotypic diversity of yeasts, lactic acid and acetic acid bacteria. Journal of Applied Microbiology, 113(6), 1428–1441. 10.1111/jam.12014

Pammi, N., Bhukya, K. K., Lunavath, R. K., & Bhukya, B. (2021). Bioprospecting of Palmyra Palm (Borassus flabellifer) Nectar: Unveiling the Probiotic and Therapeutic Potential of the Traditional Rural Drink. Frontiers in Microbiology, 12, 683996. 10.3389/fmicb.2021.683996

Pérez-Torrado, R., Rantsiou, K., Perrone, B., Navarro-Tapia, E., Querol, A., & Cocolin, L. (2017). Ecological interactions among Saccharomyces cerevisiae strains: Insight into the dominance phenomenon. Scientific Reports, 7(1), 43603. 10.1038/srep43603

Perrone, B., Giacosa, S., Rolle, L., Cocolin, L., & Rantsiou, K. (2013). Investigation of the dominance behavior of Saccharomyces cerevisiae strains during wine fermentation. International Journal of Food Microbiology. 10.1016/j.ijfoodmicro.2013.04.023

Peter, J., De Chiara, M., Friedrich, A., Yue, J.-X., Pflieger, D., Bergström, A., Sigwalt, A., Barre, B., Freel, K., Llored, A., Cruaud, C., Labadie, K., Aury, J.-M., Istace, B., Lebrigand, K., Barbry, P., Engelen, S., Lemainque, A., Wincker, P., … Schacherer, J. (2018). Genome evolution across 1,011 Saccharomyces cerevisiae isolates. Nature, 556(7701), Article 7701. 10.1038/s41586-018-0030-5

Pietrafesa, A., Capece, A., Pietrafesa, R., Bely, M., & Romano, P. (2020). Saccharomyces cerevisiae and Hanseniaspora uvarum mixed starter cultures: Influence of microbial/physical interactions on wine characteristics. Yeast, 37(11), 609–621. 10.1002/yea.3506

Pratiknjo, M. H., & Mambo, R. (2019). The Cultural Value of the Minahasa People about Liquor “Cap Tikus.” Journal of Drug and Alcohol Research, 4. 10.4303/jdar/236080

Queipo-Ortuño, M. I., Boto-Ordóñez, M., Murri, M., Gomez-Zumaquero, J. M., Clemente-Postigo, M., Estruch, R., Cardona Diaz, F., Andrés-Lacueva, C., & Tinahones, F. J. (2012). Influence of red wine polyphenols and ethanol on the gut microbiota ecology and biochemical biomarkers. The American Journal of Clinical Nutrition, 95(6), 1323–1334. 10.3945/ajcn.111.027847

Rodríguez-Lerma, G. k., Gutiérrez-Moreno, K., Cárdenas-Manríquez, M., Botello-Álvarez, E., Jiménez-Islas, H., Rico-Martínez, R., & Navarrete-Bolaños, J. l. (2011). Microbial Ecology Studies of Spontaneous Fermentation: Starter Culture Selection for Prickly Pear Wine Production. Journal of Food Science, 76(6), M346–M352. 10.1111/j.1750-3841.2011.02208.x

Santiago-Urbina, J. A., Arias-García, J. A., & Ruiz-Terán, F. (2015). Yeast species associated with spontaneous fermentation of taberna, a traditional palm wine from the southeast of Mexico. Annals of Microbiology, 65(1), Article 1. 10.1007/s13213-014-0861-8

Santiago-Urbina, J. A., & Ruíz-Terán, F. (2014). Microbiology and biochemistry of traditional palm wine produced around the world. International Food Research Journal, 21(4), 1261–1269.

Schweinberger, M. (2024). Network Analysis using R. Ladal.Edu.Au. https://ladal.edu.au/net.html

Senghoi, W., & Klangbud, W. K. (2021). Antioxidants, inhibits the growth of foodborne pathogens and reduces nitric oxide activity in LPS-stimulated RAW 264.7 cells of nipa palm vinegar. PeerJ, 9, e12151. 10.7717/peerj.12151

Silverstein, M., Bhatnagar, J. M., & Segrè, D. (2023). Metabolic complexity drives divergence in microbial communities. bioRxiv: The Preprint Server for Biology, 2023.08.03.551516. 10.1101/2023.08.03.551516

Smid, E. J., & Lacroix, C. (2013). Microbe–microbe interactions in mixed culture food fermentations. Current Opinion in Biotechnology, 24(2), 148–154. 10.1016/j.copbio.2012.11.007

Somashekaraiah, R., Shruthi, B., Deepthi, B., & Sreenivasa, M. Y. (2019). Probiotic Properties of Lactic Acid Bacteria Isolated From Neera: A Naturally Fermenting Coconut Palm Nectar. Frontiers in Microbiology, 10, 1382. 10.3389/fmicb.2019.01382

Somawiharja, Y., Wonohadidjojo, D. M., Kartikawati, M., Suniati, F. R. T., & Purnomo, H. (2018). Indigenous technology of tapping, collecting and processing of coconut (Cocos Nucifera) sap and its quality in Blitar Regency, East Java, Indonesia. Food Research, 2(4), 398–403. 10.26656/fr.2017.2(4).075

Steensels, J., Gallone, B., Voordeckers, K., & Verstrepen, K. J. (2019). Domestication of Industrial Microbes. Current Biology, 29(10), R381–R393. 10.1016/j.cub.2019.04.025

Steensels, J., & Verstrepen, K. J. (2014). Taming Wild Yeast: Potential of Conventional and Nonconventional Yeasts in Industrial Fermentations. Annual Review of Microbiology, 68(1), 61–80. 10.1146/annurev-micro-091213-113025

Stefanini, I., Dapporto, L., Legras, J.-L., Calabretta, A., Di Paola, M., De Filippo, C., Viola, R., Capretti, P., Polsinelli, M., Turillazzi, S., & Cavalieri, D. (2012). Role of social wasps in Saccharomyces cerevisiae ecology and evolution. Proceedings of the National Academy of Sciences, 109(33), 13398–13403. 10.1073/pnas.1208362109

Sumerta, I. N., Howell, K., Sudiana, I. M., & Kanti, A. (2024). Co-occurence of non Saccharomyces yeast in the natural fermentation of palm wine in Indonesia. 10.26188/25309117.v1

Tamang, J. P. (2022). “Ethno-microbiology” of ethnic Indian fermented foods and alcoholic beverages. Journal of Applied Microbiology, 133(1), 145–161. 10.1111/jam.15382

Tapsoba, F., Legras, J.-L., Savadogo, A., Dequin, S., & Traore, A. S. (2015). Diversity of Saccharomyces cerevisiae strains isolated from Borassus akeassii palm wines from Burkina Faso in comparison to other African beverages. International Journal of Food Microbiology, 211, 128–133. 10.1016/j.ijfoodmicro.2015.07.010

Tra Bi, C. Y., Amoikon, T. L. S., Kouakou, C. A., Noemie, J., Lucas, M., Grondin, C., Legras, J.-L., N’guessan, F. K., Djeni, T. N., Djè, M. K., & Casaregola, S. (2019). Genetic diversity and population structure of Saccharomyces cerevisiae strains isolated from traditional alcoholic beverages of Côte d’Ivoire. International Journal of Food Microbiology, 297, 1–10. 10.1016/j.ijfoodmicro.2019.03.001

Viljoen, B. C. (2001). The interaction between yeasts and bacteria in dairy environments. International Journal of Food Microbiology, 69(1), 37–44. 10.1016/S0168-1605(01)00570-0

Wang, C., Mas, A., & Esteve-Zarzoso, B. (2015). Interaction between Hanseniaspora uvarum and Saccharomyces cerevisiae during alcoholic fermentation. International Journal of Food Microbiology, 206, 67–74. 10.1016/j.ijfoodmicro.2015.04.022

Wijaya, L., Sumerta, I. N., Napitupulu, T. P., Kanti, A., Keim, A. P., Howell, K., & Sudiana, I. M. (2024). Cultural, nutritional and microbial perspectives of tuak, a traditional Balinese beverage. Journal of Ethnic Foods, 11(1), 4. 10.1186/s42779-024-00221-x

Xia, Q., Li, R., Zhao, S., Chen, W., Chen, H., Xin, B., Huang, Y., & Tang, M. (2011). Chemical composition changes of post-harvest coconut inflorescence sap during natural fermentation. African Journal of Biotechnology, 10(66), Article 66. 10.4314/ajb.v10i66

Yang, Y., Zhong, H., Yang, N., Xu, S., & Yang, T. (2022). Quality improvement of sweet rice wine fermented with Rhizopus delemar on key aroma compounds content, phenolic composition, and antioxidant capacity compared to Rhizopus oryzae. Journal of Food Science and Technology, 59(6), 2339–2350. 10.1007/s13197-021-05250-x

Zhou, J., Cavagnaro, T. R., De Bei, R., Nelson, T. M., Stephen, J. R., Metcalfe, A., Gilliham, M., Breen, J., Collins, C., & López, C. M. R. (2021). Wine Terroir and the Soil Bacteria: An Amplicon Sequencing–Based Assessment of the Barossa Valley and Its Sub-Regions. Frontiers in Microbiology, 11, 597944. 10.3389/fmicb.2020.597944

Ziadi, M., Gaabeb, N., Mrabet, A., & Ferchichi, A. (2014). Variation in physicochemical and microbiological characteristics of date palm sap (Phoenix dactylifera) during the tapping period in oasian ecosystem of Southern Tunisia. International Food Research Journal, 21(2).

Zongo, O., Tapsoba, F., Leray, F., Bideaux, C., Guillouet, S., Traoré, Y., & Savadogo, A. (2020). Nutritional, biochemical and microbiological composition of Borassus aethiopum Mart. Sap in Burkina Faso. Journal of Food Science and Technology, 57(2), 495–504. 10.1007/s13197-019-04078-w

Zuñiga, C., Zaramela, L., & Zengler, K. (2017). Elucidation of complexity and prediction of interactions in microbial communities. Microbial Biotechnology, 10(6), 1500–1522. 10.1111/1751-7915.12855

