## Supplementary material for "The forgotten wine: understanding the ecology and composition of palm wine fermentation": Fig S1

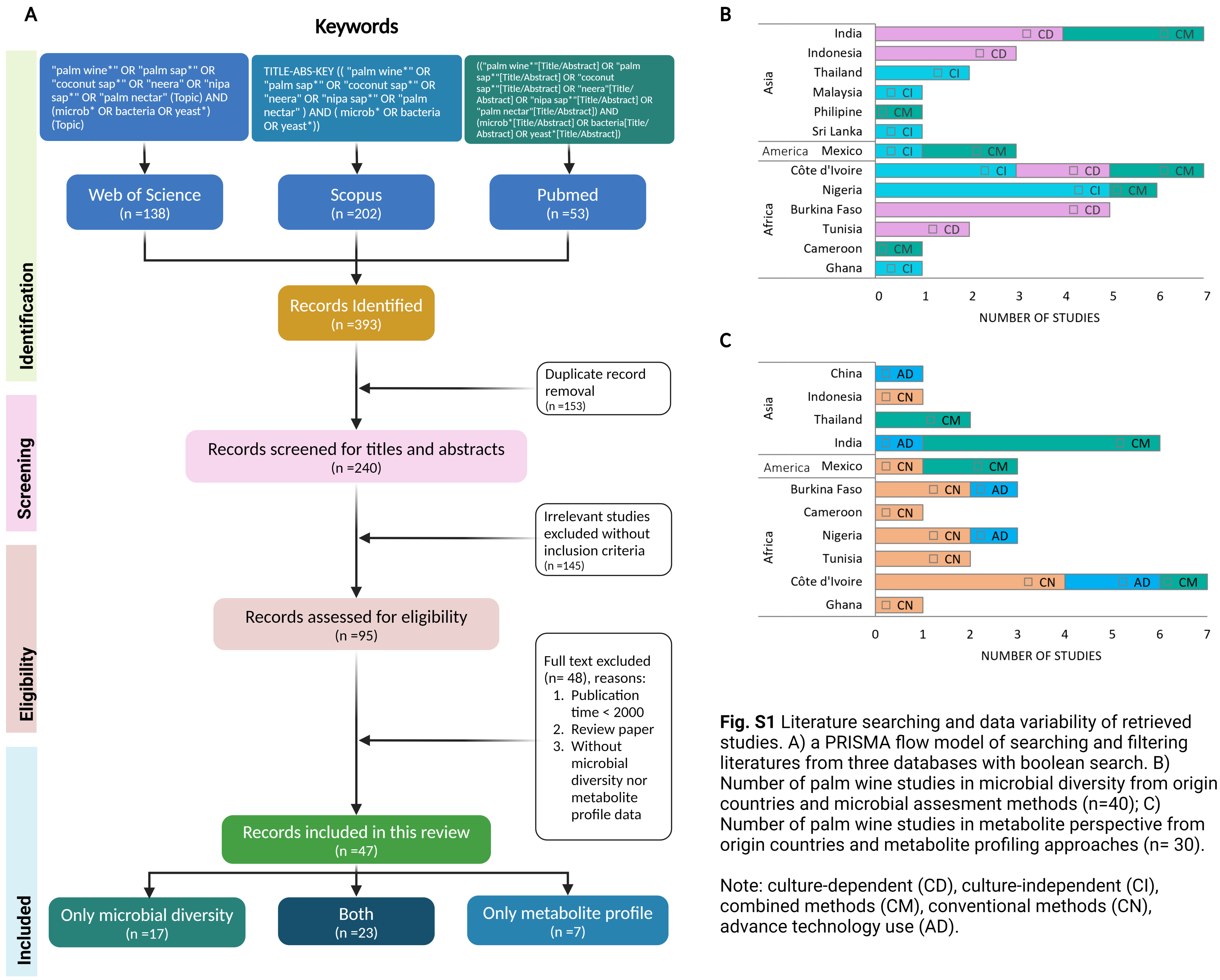


**Fig. S1** Literature searching and data variability of retrieved studies. A) a PRISMA flow model of searching and filtering literatures from three databases with boolean search. B) Number of palm wine studies in microbial diversity from origin countries and microbial assesment methods (n=40); C) Number of palm wine studies in metabolite perspective from origin countries and metabolite profiling approaches (n= 30).

Note: culture-dependent (CD), culture-independent (CI), combined methods (CM), conventional methods (CN), advance technology use (AD).
